# Artifacts in single-cell mitochondrial DNA mutation analyses misinform phylogenetic inference

**DOI:** 10.1101/2024.07.28.605517

**Authors:** Caleb A. Lareau, Michael S. Chapman, Livius Penter, Tal Nawy, Dana Pe’er, Leif S. Ludwig

## Abstract

Sequencing mitochondrial DNA (mtDNA) variants from single cells has resolved clonality and lineage in native human samples and clinical specimens. Prior work established that heteroplasmic mtDNA variants can be used to delineate clonality in hematopoiesis, but they have limited ability to reconstruct cellular phylogenies. However, a recent report by Weng *et al*. challenges the current paradigm by describing an unprecedented number of shared mtDNA variants between cells that reportedly resolve high-resolution phylogenetic trees. We re-examined the claims of Weng *et al*., and identified two major points of concern regarding this unprecedented connectedness. First, shared variants between cells are disproportionately detected in a single molecule per cell, and second, these variants are enriched 10–20-fold at the edges of mtDNA molecules, reminiscent of artifacts reported in other sequencing approaches. Further, our analyses show that pruning low support and likely artificial mtDNA variants removes nearly all of the reported phylogenetic structure. Thus, we strongly caution against using mtDNA variant calling workflows that rely on minimal evidence, including the computational pipeline introduced in Weng *et al*., as variants with high connectedness and low evidence are likely artifacts that lead to the construction of false phylogenies.

## Introduction

Cellular barcoding to trace lineages and clonality has tremendous potential to redefine our understanding of developmental and regenerative processes and to investigate disease onset mechanisms^1^. The sequencing-based identification of somatic mitochondrial DNA (mtDNA) and nuclear DNA (nDNA) mutations has revealed clonal mosaicism in nearly all human tissues, including the hematopoietic system^2–4^. It was thus with great interest that we read the report by Weng *et al*.^5^ describing Regulatory Multiomics with Deep Mitochondrial Mutation Profiling (ReDeeM), a new single-cell method reporting an unprecedented number of mtDNA variants that are shared across cells. The authors utilize this order-of-magnitude greater connectedness between cells to resolve detailed phylogenetic trees—a capability that cannot be reconciled with other contemporary results using mtDNA mutations for clonal analysis or lineage tracing^6–8^.

Longstanding work has demonstrated that whole-genome sequencing (WGS) can infer phylogenetic relationships based on somatic nDNA mutations that arise during cell division across the vast nuclear genome^9–12^. Starting from the first cell division of the fertilized egg, the lifelong acquisition of somatic nDNA variants enables the reconstruction of high-resolution phylogenetic trees^13^. In hematopoiesis, WGS of clonally expanded hematopoietic stem and progenitor cells has facilitated nuanced insights into the clonal and phylogenetic architecture of blood production across biological settings from steady-state hematopoiesis to gene therapy^9–12^. As WGS-based approaches recover mtDNA sequence at >1,000-fold coverage (due to the high copy number of mtDNA relative to nDNA), it is possible to directly assess the concordance of somatic events in both genomes and the potential for phylogenetic reconstruction from mitochondrial variants. Despite the sensitivity of calling mtDNA variants at ∼1% allele frequency, low-frequency mtDNA variants were shown to be poor markers of phylogeny compared to ground-truth nDNA data^6^. This is expected based on fundamental attributes of mitochondrial genetics, since (1) drift from ongoing mitochondrial turnover results in the loss of low variant allele frequency (VAF) mtDNA mutations over time, limiting their propagation to daughter cells; and (2) single-cell mtDNA sequencing studies reveal that clonally informative mtDNA variants typically have high mean VAF among positive cells^7,14,15^. Despite these established limitations, ReDeeM not only utilizes low-VAF mtDNA variants for phylogenetic tree construction, but it also identifies multiple large clades in young individuals covering 1–10% of total cells, whereas clonal expansions (>1% of cells) are considered extremely rare before the age of 40–50 years^10,11,16^. In this sense, the unique biological claims from ReDeeM are supported by mutations that have minimal phylogenetic signal in gold-standard contexts^6^.

Given the substantial discrepancies between ReDeeM and previous work, we revisited the methodology and results reported by Weng et al.^5^. Upon reanalysis, we found that 87.8–99.9% of connections between cells are established by variants only identified in one mtDNA molecule per cell, the minimal possible evidence. Further, these variants (or reference mismatches) disproportionately occur at the edges of sequencing reads. Such position-biased variants have previously been reported to be predominantly artifactual in other sequencing technologies^17–19^. Most importantly, removing these low-support, position-biased variants vastly depleted connectedness, leading to the collapse of the reported phylogenetic structure. Together, our reanalyses strongly caution against the analytical workflow introduced by ReDeeM, including the variant calling heuristics, tree-building analyses, and connectedness inferences.

## Results

The ReDeeM method introduces a new experimental protocol and computational workflow for analyzing mtDNA mutations from single-cell multi-omics data. Experimentally, ReDeeM enriches mtDNA from 10x Genomics single-cell multi-omic sequencing libraries and applies deep sequencing, resulting in ∼5 PCR copies (i.e., individual sequencing reads) per mtDNA molecule; a >3-fold increase in coverage compared to prior methods^5^. Combined with mtDNA variant consensus calling, the technological advances in this method depletes sequencing errors, which have been a source of false-positive heteroplasmy in mtDNA analyses^6–8^. The enrichment and consensus calling steps are important advances for higher-quality inferences surrounding mtDNA-based lineage tracing.

Although the experimental innovation in the ReDeeM method presents a useful method, the authors employ a new bioinformatics workflow with insufficient orthogonal validation. For example, the ReDeeM workflow requires only one mtDNA molecule to call a mutation in a cell (i.e., threshold >0) despite sequencing these libraries beyond saturation. This is an unusual choice, since heteroplasmic allele frequency thresholds such as ≥10% VAF are customarily used to account for (1) the multi-copy nature of the mitochondrial genome (∼150–600 copies/cell in peripheral blood^20^) and (2) the likelihood that low VAF variants undergo mitochondrial turnover and loss. For example, a variant supported by a single mtDNA molecule would only propagate in one daughter cell and disappear from the other daughter cell lineage during cell division, limiting clonal or phylogenetic inference. Likewise, the unequal segregation of mitochondria during cell division means that variants present in 2 or 3 copies have a much higher chance of being lost to cellular lineage branches during division than those at high VAFs.

We reassessed the hematopoietic cell-by-variant count matrices of the reported four different profiled donors, and observed that despite saturated sequencing, 75.4–85.1% of mutation calls within individual cells were supported by only one sequenced mtDNA molecule, representing minimal evidence (**Extended Data Fig. 1a**). Increasing the requirement to two molecules per cell causes a marked loss (87.8–99.9%) in connectedness in all four donors (**Fig. 1a,b**; **Extended Data Fig. 1b**; **Methods**), indicating the choice of binarizing at the unusual threshold of >0 is critical for the results in this work.

**Figure 1.**
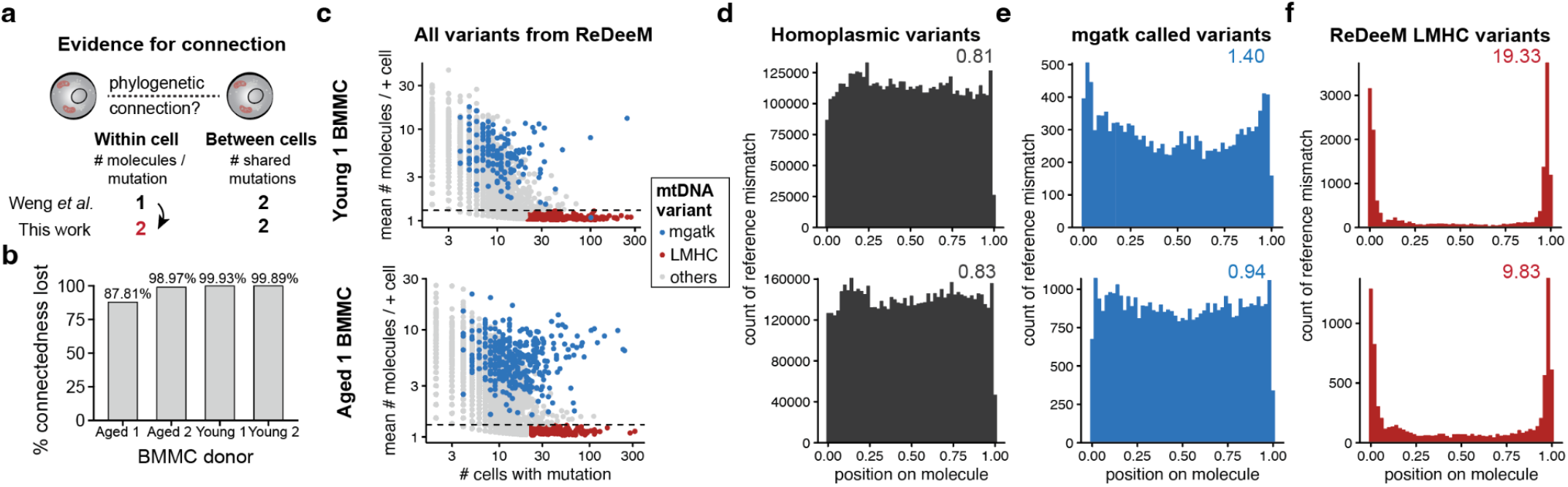
ReDeeM-based cell-cell connectivity is driven by low-mean frequency but highly abundant variants of an irregular nature. **(a)** Schematic of key parameters, including within cell (left) and between cell (right) values required to establish cell-cell connectedness. **(b)** Summary of % connectedness lost for four donors profiled by ReDeeM after increasing the required number of mtDNA molecules per mutation per cell from 1 to 2 mutations to establish connectedness between two cells. BMMC, bone marrow mononuclear cells. **(c)** Annotation of all variants called using the ReDeeM workflow. Individual variants (dots) are colored by overlap with mgatk (blue) or low mean, high connectedness (LMHC) variants (red). **(d)** Meta pileup of the position of the mismatches for all homoplasmic variants for two donors (“Young 1” - top; “Aged 1” - bottom”). Molecule fragment sizes were normalized and show the relative position of the variant, with 0.50 corresponding to the middle of the molecule and 0.00 and 1.00 to the respective ends. **(e)** Same as in (d), but for variants called by mgatk. **(f)** Same as in (d), but for the LMHC variants called only by ReDeeM. For d–f, values above each plot represent the fold enrichment of mismatches at the edges of molecules (**Methods**).

Additionally, the authors require that a minimum of two mtDNA variants be shared to establish connectedness between two cells. However, a common practice in malignancy^21,22^ and mtDNA tracing^3,7,15^ contexts is to consider single variants shared between cells, as they are clonally informative. That said, we emphasize that a single variant may be sufficient to establish clonal connectedness when the underlying variant calls are stringent and supported by multiple molecules supporting the mutation. However, the ReDeeM calls require only 1 mutated molecule per cell. Relaxing the requirement to a single shared mtDNA variant inexplicably increased the connectedness between cells reported by ReDeeM, to an average of 231 connections per cell (150–382 cells per donor; **Extended Data Fig. 1c**). This corresponds to each cell sharing connections with 2.0–4.2% of sampled cells, which would reflect large clonal expansions that contradict what has been typically observed in young hematopoietic donors^10,11^. Together, the values chosen for these two parameters (**Fig. 1a**; **Extended Data Fig. 1b–c**) deviate from extensive prior work, and minimal modifications to the chosen thresholds result in starkly different conclusions.

To resolve the mtDNA mutations responsible for this unusual connectedness, we aggregated the number of cells with the mutation observed as well as the per-variant sequencing support, defined as the count of endogenous unique molecular identifiers^5^ (eUMIs) supporting the variant per cell (**Methods**). This analysis identified a set of mtDNA variants, most of which are supported by one mtDNA molecule (mean per positive cell ≈1; threshold at 1.3), that are present in all donors and overlap in dozens to hundreds of cells. We term these “low-mean and high-connectedness” (LMHC) variants (**Fig. 1c**; **Extended Data Fig. 1d**; **Methods**). Although this set comprises only 3.9–7.1% of all ReDeeM variants, their presence in many cells disproportionately impacts cellular connectedness, a function of the number of cells squared. Critically, as low-VAF mtDNA mutations are susceptible to being lost^23^, they are unreliable for confident phylogenetic reconstruction, consistent with results from WGS reanalysis^6^. Conversely, more established variant callers, including mgatk^7^, MQuad^24^, and GATK/Mutect2^25^ require stricter thresholds per individual mtDNA variant and do not identify this set of variants (**Fig. 1c**; **Extended Data Fig. 1d,e**; **Methods**).

As other methods did not identify this LMHC variant population, we sought to examine alternative explanations that may underlie this result from ReDeeM. Specifically, prior work on variant detection in next-generation sequencing data identified artifacts at the edges of sequencing reads^17–19^, including from duplex sequencing technologies with similar consensus error correction^26^. Thus, we examined the distribution of the mismatched reference base calls underlying mutations identified by ReDeeM (**Methods**). As a control analysis, we examined the aggregate distribution of homoplasmic mutations of germline origin, verifying reference mismatches evenly across the length of molecules (**Fig. 1d**). This is consistent with the expectation that Tn5 tagmentation generates fragments randomly with respect to variants in the mtDNA genome with minor biases from Tn5 sequence preference. Similarly, heteroplasmic somatic variants identified by mgatk^7^ and MQuad^24^ are distributed relatively evenly (**Fig. 1e**; **Extended Data Fig. 1d,e**), even though their variant calling heuristics do not explicitly consider mismatch position. In sharp contrast, the aggregated ReDeeM-derived LMHC variants are ∼10–20x more abundant at the edges of mtDNA molecules than at their center for all four donors (**Fig. 1f**; **Extended Data Fig. 1d**; **Methods**). Further, nine additional donors profiled via a hashed ReDeeM workflow are also enriched 6.1–31.8-fold for reference mismatches at the edges of molecules (**Extended Data Fig. 2a**), suggesting pervasive edge bias in all ReDeeM libraries for 13 total donors. Together, our analyses show that the ReDeeM pipeline is unique in identifying a set of variants supported by only one detected mtDNA molecule per cell, and these variants exhibit strong positional bias resembling reported artifacts^17–19^.

Notably, other sequencing technologies that similarly use consensus calling explicitly filter edge artifacts or trim the edges of reads^26,27^ though edge trimming tools were inexplicably not utilized in the ReDeeM workflow. For example, a recent report of duplex sequencing for mtDNA mutations in conplastic mice strains reported a residual error rate of ∼2×10^−8^ nucleotides after consensus calling^28^. This report further noted an artificial enrichment of reference mismatches at the 5’ end of reads, which were subsequently trimmed to exclude erroneous variant calling^28^. Noting the pervasive accumulation of edge mismatches, we observed a similar edge-position bias in LMHC variants in the original ATAC library as well as independent single-cell and bulk ATAC-seq libraries, from both mtDNA- and nuclear-derived accessible chromatin fragments (**Extended Data Fig. 2b–d**). These analyses indicate that spurious mismatches at the edges of sequenced DNA molecules are a technical feature of ATAC-seq data, which are not removed by the consensus calling approach used in ReDeeM (**Methods**).

To quantify this edge position bias at the variant level, we developed a heuristic that is inspired by the gatk ReadPosRankSumTest^29^, but is compatible with the ReDeeM data structure (**Methods**). Specifically, we use a Kolmogorov-Smirnov statistic to assess where reference mismatches occur on the molecule. To conform with the ReDeem data structure, we specified an expected background distribution of uniform occurrence of the mismatch across the molecule (rather than the distribution of the reference match alleles in gatk^29^). Application of this statistic to all heteroplasmic and homoplasmic mutations revealed an empirical cutoff of 0.35 that did not flag any homoplasmic mtDNA variants (i.e., true variants; **Extended Data Fig. 3a; Methods**), which we used for interpreting heteroplasmic mutations. Only 1.5–3.8% of heteroplasmic mtDNA variants called by mgatk exceed this threshold, whereas 70–88% of ReDeeM LMHC variants and 23–32% of all other ReDeeM variants deviate from the expected uniform distribution (**Extended Data Fig. 3a**). Analysis of all ReDeeM-identified variants showed that this edge bias is inexplicably most pronounced when exactly one or two mtDNA molecules support the mutation in a cell, whereas variants with 3 or more supporting molecules exhibit no such bias (**Extended Data Fig. 3b**). In addition, ReDeeM-called mismatches are 7–24-fold enriched per donor for transversions when supported by only 1 compared to 3+ eUMIs (**Extended Data Fig. 3c**). As transition mutations are much more common in mtDNA from hematopoietic cells (typically 90–95%)^3,7^, the underlying nucleotide composition of ReDeeM variant calls with low eUMI support reinforces that edge-enriched variants are likely artificial.

Together, these analyses indicate that the unprecedented connectedness reported by ReDeeM is disproportionately due to mtDNA variants supported by only one molecule per positive cell. These low-support variants called by the ReDeeM pipeline and not other workflows display pronounced irregularities that are more readily explained by technical artifacts than biological signal. To assess the downstream impact of these low support and likely artificial variants, we constructed dendrograms for the bone marrow mononuclear cell data for four donors using the weighted Jaccard distance and neighbor-joining algorithms proposed by Weng *et al*.^5^ (***Methods***). Trees that we reconstructed using the full ReDeeM variant set closely resemble the hierarchical structure in the original report^5^ (**Fig. 2a**). We next assessed the degree of phylogenetic signal that is retained using variant calling thresholds with greater biological confidence. By requiring 2+ molecules of support per mtDNA variant, the revised trees exhibited a starkly dissimilar topology (**Fig. 2b**). To quantify the degree by which the two trees share phylogenetic signal, we used a most recent common ancestor (MRCA) trio analysis^3^ (**Methods**), finding near-random relationships between the original tree and the one requiring 2+ eUMIs (33.8–39.2%; **Fig. 2c**, grey; random is 33.3%). Conversely, variants with exactly 1 eUMI retain a substantial fraction of MRCA trio concordance (64.7–68.8%; **Fig. 2c**, black), underscoring the divergent information content of lowly abundant and likely artificial mtDNA variant calls as compared to more bona fide mtDNA variants. Although a CRISPR-based lineage-tracing mouse model was used to validate phylogenetic inferences based on ReDeeM variants, our examination of these data revealed at least five experimental or computational differences from the human ReDeeM pipeline, none of which were documented in the original report by Weng et al.^5^ (**Methods**). Despite the fundamental differences in data composition and workflows, an accumulation of LMHC variants and edge bias of supporting molecules is also present in the mouse data (**Extended Data Fig. 4**). Other purported validations reported by the authors, such as the clonal *k-*nearest neighbor (*k-*NN) analyses, had independent logical inconsistencies that one cannot consider rigorous validation of the ReDeeM methodology (**Methods; Supplemental Note**). Finally, we note that additional sources of confounding may still limit the interpretation of these ReDeeM data, such as ambient mtDNA, which would also drive spurious phylogenetic inferences **(Supplemental Note**).

**Figure 2.**
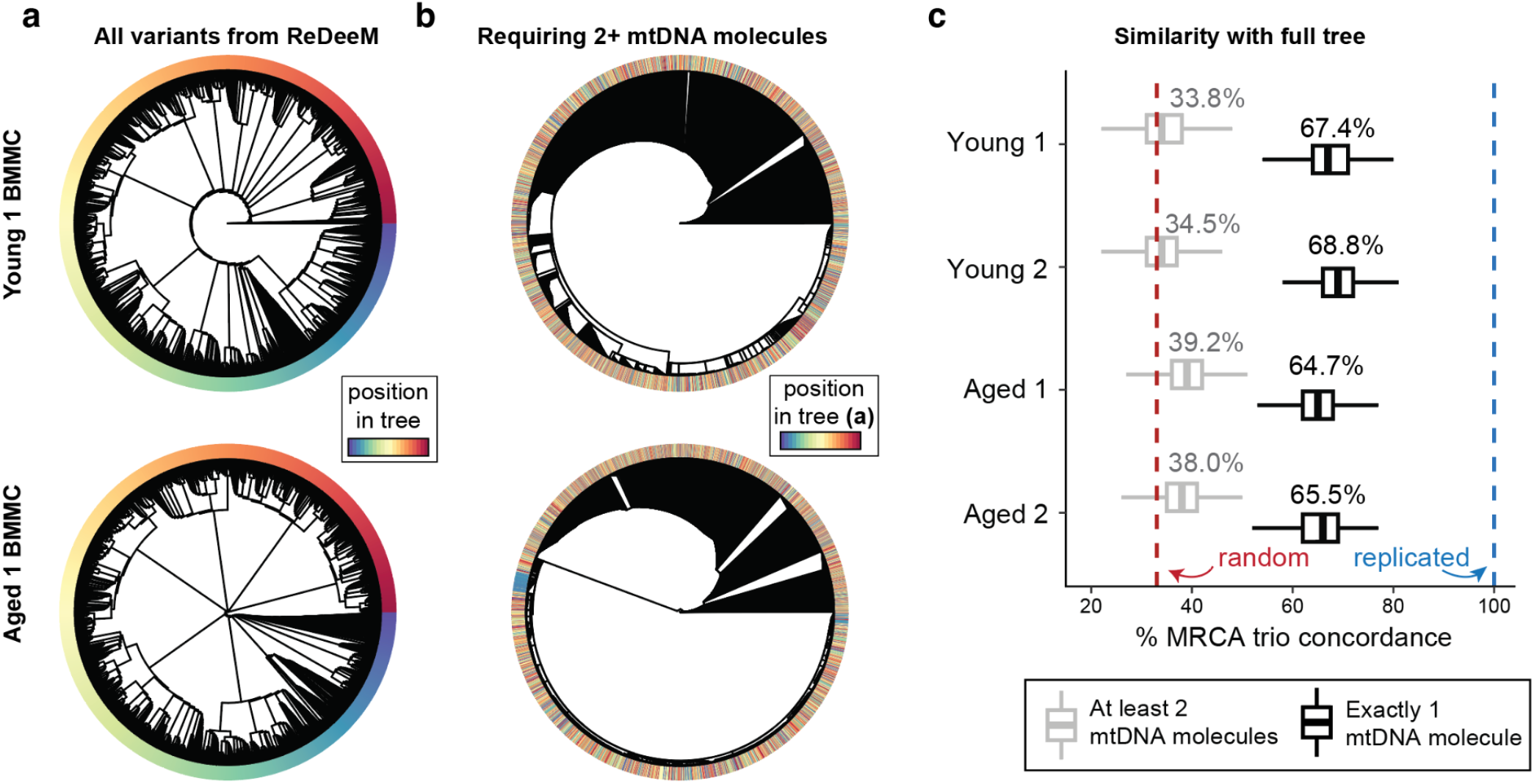
Phylogenetic tree inferences are primarily driven by variants supported by only one molecule. **(a**,**b)** Phylogenetic trees of BMMCs from Young 1 and Aged 1 individuals were constructed using (**a**) all variants from the ReDeeM pipeline or (**b**) only variants that are supported by 2+ mtDNA molecules. The color bar marks the relative position of cells in the topology in (a), showing substantial reshuffling of cells following removal of cell/variant calls with exactly one molecule of support in (b). **(c)** Quantification of tree similarity between (a) and (b) using trio analyses to determine most recent common ancestors (MRCA), when considering variants with 2+ mtDNA molecules of support (grey), compared to only considering variants supported by exactly 1 mtDNA molecule (black). Box plots represent 100 bootstraps each of 1,000 trios that were compared using the tree in (a) as the “truth” and the tree in (b) as the “query”. MRCA trio analyses would yield a mean of 33.3% for completely random trees (dotted red line).

## Conclusions

Some innovations introduced by Weng *et al*.^5^, including the targeted capture, deep sequencing, and consensus calling of variants on mtDNA molecules, hold promise for establishing more sensitive single-cell mtDNA genotyping. However, our reanalysis reveals the existence of extensive technical artifacts upstream of library generation and sequence alignment, which cannot be corrected by PCR consensus calling and demand greater supporting evidence for confident mtDNA variant calls^17–19^. Most mtDNA variants supported by exactly one molecule display a likely artificial edge-biased distribution, undermining their utility in downstream analyses, including phylogenetic reconstruction. These low-support artifacts, when called as variants, greatly exaggerate variant abundance and strongly distort the inference of cell-cell connectivity and phylogeny, which is effectively random compared to variant-calling heuristics requiring stricter thresholding (**Fig. 2**). Independent of these major technical concerns, low VAF mtDNA variants are most susceptible to being only propagated in one of the daughter cells, thus generally undermining their biological utility for any type of phylogenetic or clonal inference.

Independent single-cell multi-omics approaches^3,7^ and WGS of single-cell-derived hematopoietic colonies^6,30^ (considered the gold standard and highest resolution method for retrospective phylogenetic inference of human cells with sensitivity to detect mtDNA variants at <1% VAF) further indicate that somatic mtDNA mutations provide limited phylogenetic, rather than clonal, information in hematopoiesis. Specifically, WGS demonstrates that most mtDNA mutations accumulate throughout life, rather than during the relatively brief period of development when the HSC pool is specified. The highly connected phylogenetic tree proposed by Weng et al.^5^ would require hundreds of mutations to barcode the development of individual HSCs, which is inconsistent with other inferences of mtDNA variant accumulation^3,6,23^. We therefore strongly caution against the bioinformatic methods proposed in the ReDeeM workflow, particularly the reconstruction of human hematopoietic cell phylogenies via mtDNA mutations, noting the highly unusual and almost certainly artificial support underlying these unique claims. In particular, we stress that other widely-used variant calling workflows, including those using consensus calling^28^, would discard edge variants to exclude erroneous variant calls. Conversely, ReDeeM provides no justification or proper benchmarking to warrant their inclusion. Future work will be required to reconcile the advances presented in Weng et al.^5^ with heuristics that mitigate these artifacts. Given the turnover and (asymmetric) distribution of mitochondria and genomes within during cell division^31^, an improved understanding of mtDNA copy number and somatic mtDNA heteroplasmy dynamics will be required to bolster mtDNA-based clonal and lineage tracing and to assert confident retrospective inference in the hematopoietic system and other human tissues.

## Author contributions

C.A.L. led bioinformatics analyses with critical input and guidance from M.S.C., L.P., T.N., D.P., and L.S.L. All authors participated in the drafting and revising of the manuscript.

## Data availability

Data from this work was downloaded from **GSE219015** which was previously published.^5^

## Code availability

Custom analysis code used to reproduce all inferences in this response is available at: https://github.com/caleblareau/redeem-reanalysis-reproducibility.

---

**Extended Data Figure 1.**
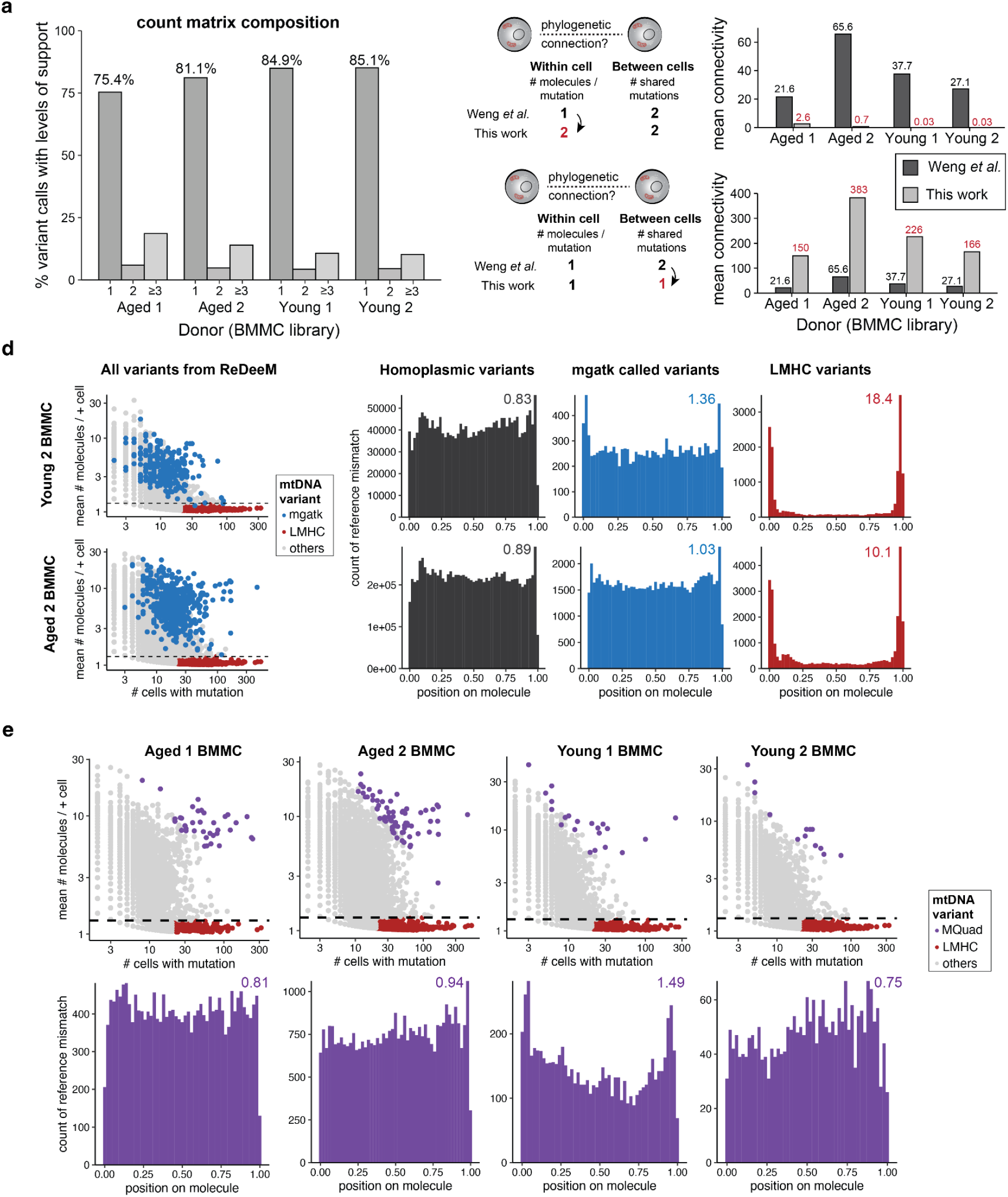
The majority of ReDeeM-called variants have low support and are likely artificial. **(a)** Summary of support per variant per cell, stratified by donor, represented as the percent of non-zero entries in the cell-by-variant ReDeeM matrix. The percent of variants with exactly 1, 2 or ≥3 molecules of support per cell appears above the corresponding bars. **(b**,**c)** Connectedness analyses, including key parameter adjustments (left) and the effects on mean connectivity (right). **(d)** Same analyses as in **Fig. 1**, but for the two other donors profiled by Weng et al., showing similar positional bias in LMHC variants. **(e)** Annotation of all variants called using the ReDeeM workflow, including variants called by MQuad (purple) for four donors (top). Meta pileup of the position of the mismatches for variants called by MQuad (bottom). The values above each plot represent the fold enrichment of mismatches at the edges of molecules.

**Extended Data Figure 2.**
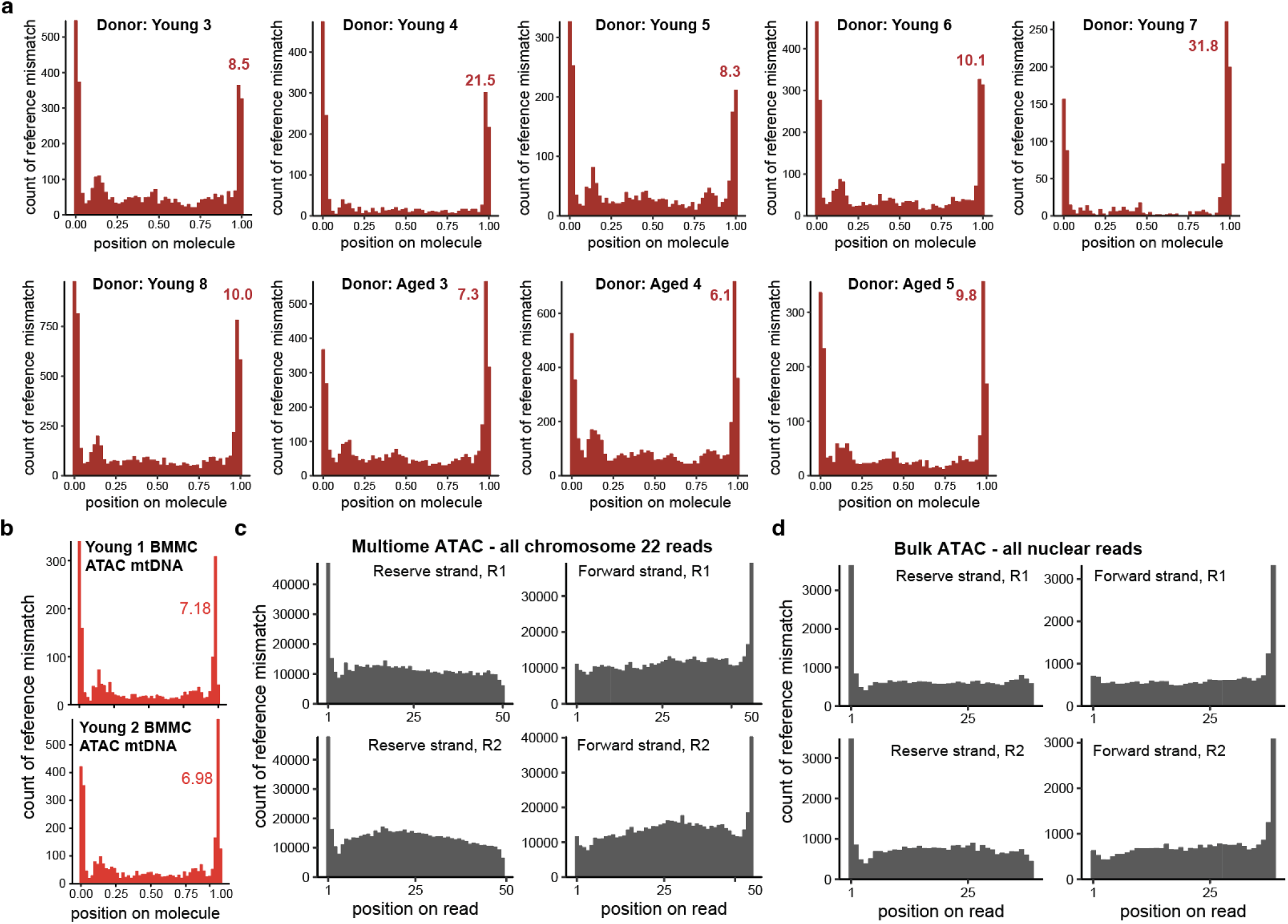
Mismatch position bias is present throughout ATAC-seq-derived molecules. **(a)** Pileup of LMHC variants for 9 additional donors from the ReDeeM hashing experiments. Values in each panel represent the fold enrichment of mismatches at the edges of molecules relative to the center. **(b)** Pileup of LMHC variants from **Fig. 1c** and **Extended Data Fig. 1d** from the multiome ATAC-seq library. Values in each panel represent the fold enrichment of mismatches at the edges of molecules. **(c)** Pileup of all positions containing a mismatch or reads mapping to the reverse strand or forward strand of chromosome 22, split by the sequencing read, from a publicly available single-nucleus peripheral blood mononuclear cell multiome experiment (**Methods**). **(d)** Same as in (c) but for a bulk ATAC-seq experiment^3^ summarizing all nuclear chromosomes. Note that no variant calling was performed for panels c and d, such that pileups reflect all mismatches from the reference observed in the library.

**Extended Data Figure 3.**
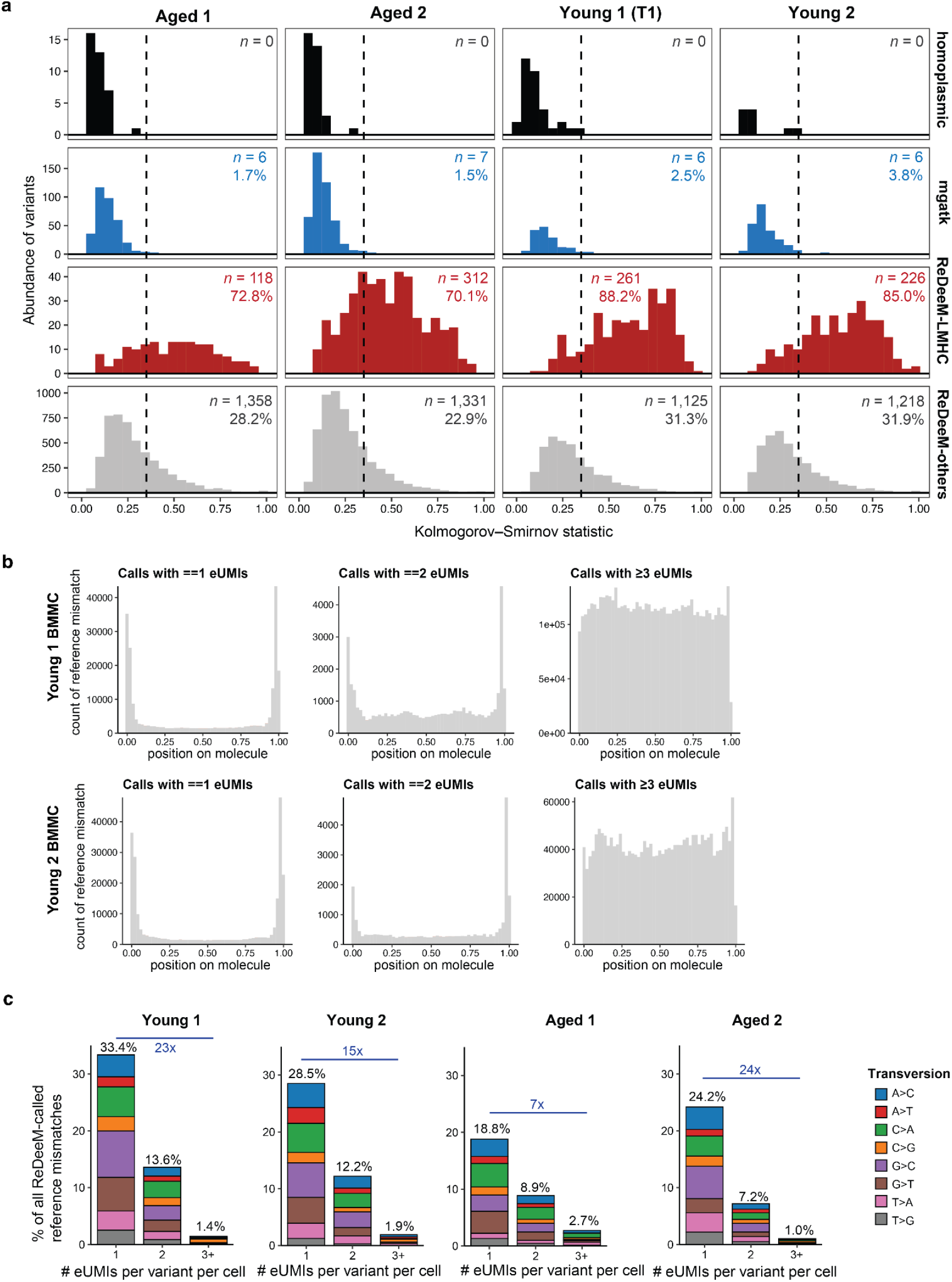
A substantial set of ReDeeM variants exhibit artifactual features. **(a)** Kolmogorov-Smirnov (KS) statistics for variants identified via different methods. The number and percentage of variants with a KS statistic > 0.35 (dotted line) are reported per panel. **(b)** Pileup of all ReDeeM-called heteroplasmic variants, stratified by the amount of support (i.e., eUMIs per variant per cell). Each variant is represented in two or more panels per donor. **(c)** The composition of nucleotide changes at the edges of molecules, stratified by the number of eUMIs detected for the mutation. Percentages represent the sums of all transversions; blue numbers represent the fold enrichment of transversions between 1 eUMI and 3+ eUMI categories for each donor.

**Extended Data Figure 4.**
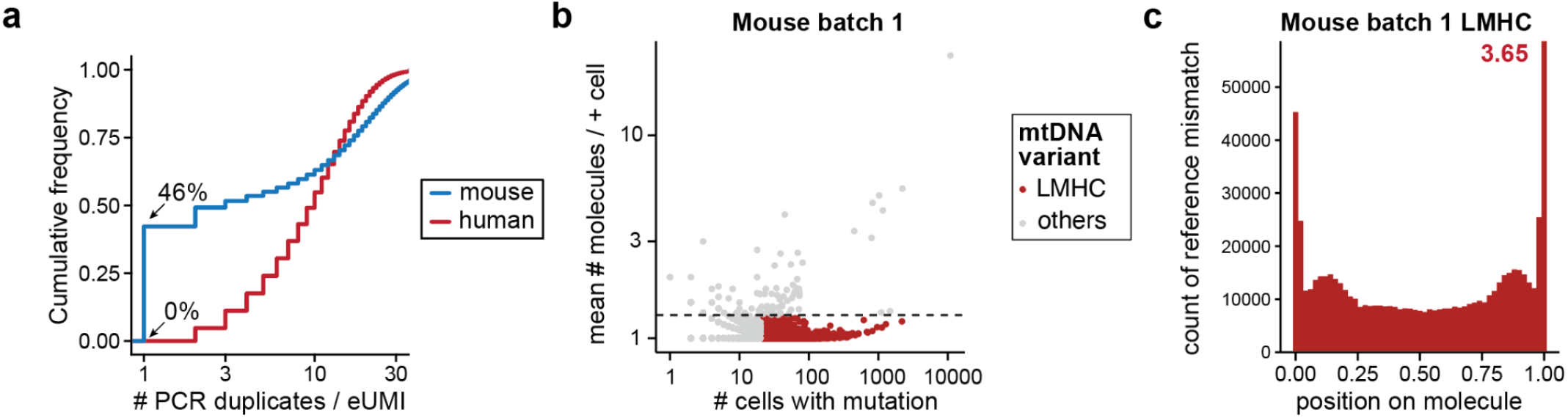
Comparison of human hematopoiesis and murine CRISPR tracing tumors. **(a)** Cumulative distribution plot comparing the number of PCR duplicates per endogenous unique molecular identifier (eUMI) in the Young 1 human bone marrow dataset (human) and a mouse Kras;Trp53 (KP) tumor model. Highlighted percentages represent the number of eUMIs with exactly 1 PCR duplicate. **(b)** Annotation of low mean, high connectedness (LMHC) variants in the mouse KP tumor validation data. **(c)** Summary of reference mismatch positions in the murine data for the LMHCs in (b).

## Methods (Supplementary Information)

### Reanalysis of ReDeeM samples

Thirteen single-cell multi-omic libraries from four human donors (Young 1, Young 2, Aged 1, and Aged 2) profiled by ReDeeM comprised the available data when the manuscript was originally made available online (January 27, 2024). We focused on libraries profiled from bone marrow mononuclear cells (BMMC) for all four donors as other populations (e.g., hematopoietic stem and progenitor cells) were not profiled consistently. We selected the BMMC samples for each donor (the first time point for the first donor), as these presented the major datasets benchmarking ReDeeM in the original report. However, all thirteen datasets were benchmarked and showed a similar abundance of LMHC variants, molecular position bias rates, and reduced connectivity per donor as shown for the BMMCs. Low mean and high connectedness (LMHC) variants are defined by having fewer than 1.3 mean molecules per positive cell, and at least 20 cells per sample (red variants in **Fig. 1c** and **Extended Data Fig. 1d**).

### ReDeeM connectivity analyses

Based on parameters used in the “BuildTree.R” function in the redeemR package (v.1.0.0; released December 2, 2023) and “Note-2” from the ReDeeM-reproducibility repository (updated February 25, 2024), we excluded mtDNA variants that are highly abundant across all four donors: 3109T>C, 5764C>T, 309T>C, and 310T>C, which are notoriously hard to genotype because of the low complexity of the genome near these positions. Of note, including these four variants would have resulted in a more extreme before- and after-impact on connectedness. Further, for the Aged 2 donor, we removed two variants (72T>C and 10810T>C) that were classified by ReDeeM to be heteroplasmic but had unusual distributions (96% and 80% heteroplasmy respectively among positive cells). For the other three donors, no additional variants were removed upstream.

Aside from these exclusions, we used the default “Sensitive” parameters for all ReDeeM variant calls. The same methodology to compute the connectedness introduced in the ReDeeM manuscript was used to compute the baseline connectedness, and our median values were the same as what was originally reported. In one line of code, we modify the binarization to be >= 2 rather than >= 1 and then report the change in the sum of off-diagonal connectedness (**Fig. 1b**). A similar change in a single line of code assessed the curious choice of requiring 2+ mutations to establish connections between cells. Relaxing this parameter to quantify only 1 shared mutation (which is more consistent with prior results of mutation sharing from other mtDNA data^3,7^) resulted in connections per cell with 2.0–4.2% of other cells in the experiment, reflecting a large degree of clonal expansions inconsistent with measures of clonality, in particular at a younger age in human hematopoiesis^10,11^. In light of our characterization of the likely artifacts underlying the ReDeeM data, we interpret this parameter choice as a means of obscuring high connectedness driven by largely artificial mutations.

### Comparison with other bioinformatics methods

Here, we summarize the ReDeeM bioinformatics pipeline (hereafter, ReDeeM-V as reported^5^) with other state-of-the-art mtDNA variant calling tools. We emphasize that none of the other three tools show a substantial (i.e., >2-fold) edge enrichment for called variants, nor consider a reference mismatch on a single molecule as sufficient evidence for a cell to have an mtDNA variant.

#### mgatk

A set of high-quality variants is determined by aggregating information across cells^7^ to create per-variant metrics. Notably, subclonal analyses with mgatk require at least 3 cells that are “confidently detected”, meaning that at least 2 reads span the reference mismatch from both orientations (for a minimum of 12 sequencing reads to be called a mutation). Thus, most of the LMHC variants identified in the ReDeeM workflow provide insufficient per-cell evidence for identification via mgatk. For comparisons with the ReDeeM results, we executed mgatk on the CellRanger-Arc.bam files from the ATAC-seq library (as previously demonstrated^14^). Thus, the mgatk call set reflects the variant calls without the mtDNA enrichment and only the mtDNA that is present from the transposition reaction. The choice of parameters for variant filtering was the same for mgatk as used in the ReDeeM paper^5^ and our past work on the multiome kit^14^.

#### Mquad

MQuad^24^ uses a two-component mixture model to identify variants with sufficient evidence of a biological signal in the overall heteroplasmy matrix. Specifically, a Bayesian Information Criterion (BIC) score is used to determine which variants should be included in downstream analyses. We ran MQuad with the default parameters on “AD” and “DP” matrices as input. We adopted the output of the ReDeeM data structure to generate these variants to utilize only the processed ReDeeM data structure. Consistent with our experience in other datasets, MQuad^24^ returns a stringent but likely incomplete set of mtDNA variants, as it requires substantial evidence for variant calling (**Extended Data Fig. 1e**). As MQuad enriches for high-confidence variants, the pileup profile was largely void of edge bias, and LMHC variants were excluded due to lack of read-level evidence. The set of variants called by MQuad were those that exceeded the BIC threshold based on the default knee plot, but we note that more lenient thresholds (e.g., top 500 variants) were similarly depleted of edge-biased variants.

#### ReDeeM-V

For the four donors deeply analyzed in the ReDeeM manuscript, the authors require the “S” or “Stringent” mode for analysis, requiring at least 2 cells to have detected 1 or more eUMIs and at least 1 cell to have 2+ eUMIs for the variant to be called. Thus, a minimum of 3 eUMIs out of hundreds of millions of sequencing reads is the minimal evidence required for a variant to be considered in downstream analyses, noting that additional post-hoc filters are applied to further filter mutations that are disproportionately strand-biased. The authors indicate that these very lenient thresholds can be applied because of the error correction steps in eUMI consensus calling. For clarity, we agree with the suggested workflows to minimize sequencing errors, but we stress that other artifacts during library preparation would not be corrected, including those near the edges of sequencing fragments.

#### GATK / Mutect2

Originally designed for bulk sequencing datasets, conventional callers such as Mutect2 (part of the GATK workflow) currently operate on each cell individually; unlike the other tested methods, they do not allow for information sharing between cells. Under this workflow, mutations are called probabilistically, but would always require more than one PCR de-duplicated read for variant calling (in our experience, 3 or more are typically required). Thus, the vast majority of LMHC variants in single cells would not be identified using the conventional variant calling approach of Mutect2, but likely at the expense of real mutations that can be confidently identified in other cells. To enhance sensitivity, one approach is to identify mtDNA variants in individual cells using Mutect2 and then summarize them as a union across cells to generate a counts matrix, as was recently utilized to analyze plate-based single-cell DNA sequencing data^25^. However, as an additional filtering mechanism, cells with variants were required to have at least two PCR deduplicated reads for calling a potential variant in individual cells, which would result in the same magnitude of change as we show in **Figs. 1** and **2**.

### Analyses of variant positional bias

For meta-variant pileups, the relative position of each variant was determined by the genomic coordinate of the mutation of the left and right transposition events. The numeric edge bias value was computed as the ratio of the average of end-most regions on either side of the molecule (edge 5% on either side) over the average of the remaining bins throughout the molecule.

Inspired by the gatk ReadPosRankSumTest^29^, we utilize a Kolmogorov-Smirnov (KS) statistic to identify variants with skewed distributions of reference mismatch support. The key difference between our convention and the gatk test is the different definitions of the null distribution. Whereas gatk defines the position along the molecule for the reference allele, we used a uniform [0,1] distribution as our null to comply with the processed ReDeeM data structure. As the eUMI molecules were only reported if they contained one or more mismatches, the full distribution of the reference alleles was not accessible in the processed ReDeeM data alone. Due to the large number of reference mismatches per variant, we elected to use the statistic rather than the *p*-value and found this to be a more reasonable classifier. An empirical threshold of 0.35 was selected as this value retained all homoplasmic (i.e., true positive) variants for all four donors (**Extended Data Fig. 3a**) and produced a classification of variants that met expectations visually (examples are available through the online code resources). Future work that modifies the ReDeeM data structure to allow for a null distribution of the reference allele will likely improve accuracy in filtering artificial variants with molecular position bias.

### Public data access and processing

To assess the possibility of spurious mismatches in the ATAC-seq library from the multiome kit, we downloaded a bam file summarizing a peripheral blood mononuclear cell profiling experiment from 10x Genomics (the “10k_PBMC_Multiome_nextgem_Chromium_X_atac” bam prefix). Aligned reads were parsed for the “MD” tag that reports the mismatch from the reference position. We aggregated the occurrence of reported mismatches without performing variant calling, meaning that more background will be present in the form of sequencing and other errors (**Extended Data Fig. 2c**). Similarly, we reprocessed a bulk ATAC-seq experiment (SRR7245894^3^) with analogous upstream alignment and “MD” tag parsing steps (**Extended Data Fig. 2d**). Regardless, the accumulation of mismatch positions at the edges of the fragments strongly support the notion that the artificial signal driving many variants in ReDeeM (**Fig. 1f**) is attributable to the underlying molecular biology processes in the ATAC-seq chemistry.

### Mouse ReDeeM data

Weng *et al*. utilized a CRISPR lineage recording Kras;Trp53(KP)-driven lung adenocarcinoma tumor model to validate phylogenetic inference. They co-detected mtDNA alongside the CRISPR evolving barcode and concluded that the reported concordance between the CRISPR barcodes and mtDNA mutations validates the ReDeeM method. Conceptually, these analyses did not quantify lineage fidelity at the single-cell level (i.e., the phylogenetic relationship of individual cells), which is implied by the creation of single-cell dendrograms. Instead, the mtDNA variant data was used to predict the clonal groupings of the evolved CRISPR barcode. Thus, we suggest that the more appropriate interpretation of the mouse data is restricted to the clonal level (where mtDNA tracing has been previously established^3^) rather than fine-grained, single-cell phylogenies as reported.

Regardless of the design and interpretation of the experiment, we note there were multiple discrepancies between how the authors reported the murine analyses compared to what was performed, based on public metadata from the processed files. Therefore, we separately caution against the interpretation of this experiment as the data analyses differ from the reported human workflow in at least five critical areas:

1. *Library separation –* The authors did not provide a separate mtDNA-specific library as part of the data sharing (as of July 15, 2024) and instead analyzed mtDNA and ATAC-seq from the same sequencing library.
2. *Sequencing saturation –* As no dedicated mtDNA library was isolated, the saturation to 8 PCR duplicates per eUMI could not be achieved as reported for the human samples (**Extended Data Fig. 4**).
3. *ReDeeM-V threshold* – In contrast to the human samples that used a ReDeeM-V/ReDeeM-R pipeline with stringent (“S”) setting for downstream analyses, the murine data used the total (“T”) setting. Though the overall impact of this parameter choice is not clear, the combination of this new threshold and the limited sequencing saturation leads to a fundamentally different distribution of PCR duplicates per eUMI (**Extended Data Fig. 4a**), likely resulting in far more sequencing error than the human datasets.
4. *Alignment software* – Whereas the human ReDeeM libraries were aligned to bowtie2, the murine data was processed with CellRanger-ATAC, which uses bwa as an aligner. In our experience, these two aligners vary in their treatment of reference mismatches at the edges of molecules; bwa soft-clips the edges of molecules more often, which may explain why the relative enrichment at the edges of the molecules are less pronounced in the murine data.
5. *Reference genome masking* – best practices for ATAC-based mtDNA tracing include the masking of nuclear-encoded mitochondrial DNA (NUMTs) to increase reference coverage^32^, which was used for the human analyses in the ReDeeM manuscript. However, for the mouse data, this was not utilized, resulting in regions of the mtDNA genome being poorly covered and limiting inferences.

Ultimately, the impact of these parameter choices is unclear. Regardless, the presence of LMHC variants and edge bias still indicate that the artificial signal underlying the ReDeeM method is sustained in the murine data. Due to differences in choices for these experimental and bioinformatics methods, the degree of edge bias is likely somewhat minimized as sequencing errors will be more abundant in the absence of consensus calling as sequencing errors will occur throughout the molecule with a more modest enrichment at edges than we detected in the ReDeeM data. Ultimately, we emphasize that careful and complete documentation is required to conclude that the mouse data can validate the clonal or phylogenetic signals of mtDNA variants in the hematopoietic system.

### *k*-Nearest Neighbor analyses

As part of the original article’s Supplementary Note “Enumeration of multifaceted ReDeeM validations,” the first data point offered is that the major hematopoietic lineages are recapitulated in a *k-*nearest neighbors (*k*-NN) analysis. Separate and in addition to major concerns in the underlying ReDeeM workflow described throughout this manuscript, we suggest that this is an inappropriate analysis. In the “Lineage origins of hematopoietic cell types” section, the authors remove 85-90% of all called mtDNA mutations as they do not conform to desired branches of hematopoietic differentiation (computed via binomial tests). Using the remaining 10-15% of mutations that are enriched in one of the four lymphoid or myeloid/megakaryocyte/erythroid branches, the authors compute a *k-*NN graph from the filtered variant by cell matrix. By restricting the feature set to only mutations that fit this predefined expectation of hematopoietic lineage bias, the *k-*NN analysis trivially shows a greater degree of clonal relatedness within each major hematopoietic branch.

On the other hand, the excluded set of variants is conceivably enriched in multipotent stem and progenitor cells that largely equally contribute to major branches, and would thus not conform with the “expected hematopoiesis differentiation process”. However, these ‘multipotent’ variants nevertheless conform to a biological expectation of multi-lineage differentiation and would not be excluded from downstream analysis. Thus, the *k*-NN analysis can not be considered a rigorous validation of ReDeeM or underlying mtDNA variant quality; instead, this analysis reports a heavily skewed result.

### Comparison of trees

As the GitHub resources associated with the original paper loaded tree objects that were not made available (git commits on February 25, 2024; last update July 15, 2024) and that we could not reproduce with our best efforts, the trees presented in **Fig. 2** do not represent a direct comparison to the original report. Rather, we elected to compare trees using the methods described by the authors (namely, ReDeeM ‘Sensitive’ variants to compute the weighted Jaccard distance matrix and then neighbor-joining). We used the identical tree-building procedure to compare the “after”, the only difference being that we partitioned the matrices to either include variant calls with only 1 (**Fig. 2c**, black) or 2+ eUMIs (**Fig. 2c**, grey). To assess concordance with the full tree, we used a most recent common ancestor (MRCA) trio analysis^3^, which classifies trios of arbitrary cells of whether a new tree agrees on the two that are most closely related.

Boxplots represent 100 bootstrap estimates of 1,000 trios sampled randomly from the pairs of trees. Across the four donors, the MRCA concordance using variants with 2+ eUMIs of support ranged from 33.8–39.2%, a modest increase from 33.3%, reflecting minimal overlap between pairs of trees. In contrast, the trees using only 1 eUMI had higher values (64.7–68.8%), reflecting that most of the purported phylogenetic signal in the original ReDeeM trees are driven by variant calls with only 1 eUMI of support per cell. Taken together, given the genetic properties of mtDNA where 2+ mtDNA molecules must be present for valid lineage relationships, we urge strong caution when generating and interpreting phylogenetic trees from single-cell mtDNA genotyping data, whether from ReDeeM or other related technologies. More generally, our analyses caution against the use of the bioinformatics workflow introduced in the ReDeeM manuscript^5^.

As of August 25, 2024, the authors have provided a new filtering method for the ReDeeM data that explicitly remove variants that accumulate on the edges. The authors suggest that the topologies of the trees are informative for phylogenetic inferences with either method. Our evaluation of these trees that are produced with the two versions of the ReDeeM filtering yield essentially random with respect to each other (**Supplemental Note**), similar to our analyses shown in **Figure 2**.

