## Supplemental Note for "Artifacts in single-cell mitochondrial DNA mutation analyses misinform phylogenetic inference"

This document summarizes notable inconsistencies in the authors' response[1] to our original submission and preprint[2] as well as alternative explanations for data presented by the authors considered evidence of the robustness and reliability of ReDeeM.

**Context**

- We previously expressed fundamental concerns that the ReDeeM bioinformatic pipeline identifies artifactual mitochondrial DNA (mtDNA) variants, which contribute to a high degree of cell-cell connectedness and drive phylogenetic inference[2].
- In response, the original authors posted a preprint, introducing a new ReDeeM "filter -2", which removes alleged artifact variants[1]. The authors argue that the new filter -2 performs similarly to the original ReDeeM workflow[3] (filter -1), suggesting that ReDeeM produces robust results with either filtering approach, including phylogenetic inference. No recommendations are made concerning the context in which either filter should be applied, including whether filter -1 is ever appropriate to use.
- The authors provide arguments and analyses claiming that variants supported by only one molecule are informative and should be retained for phylogenetic inference.
- In their conclusion[1], the authors appear to acknowledge that these inferences may not always be "perfect" and suggest that their tool may be most suitable for "exploration".

**Results comparing ReDeeM filters -1 and -2**

We posit that the evaluation metrics used in the author's response are not appropriate for benchmarking the two ReDeeM filters. The major inconsistencies of these analyses and conclusions are:

- Quantification of the connectedness metric (introduced in the original ReDeeM publication[3]) or cell-cell connectivity (used in the preprint[1]) both show that despite removing only ~5% of (artifactual) mutations, ReDeeM filter -2 reduces the connections by ~53-99% compared to ReDeeM filter -1 (**Supplemental Fig. 1**). We again emphasize that not all mutations contribute to phylogenetic reconstruction equally, and the focus on the frequency of filtered mutations rather than the resulting connectivity does not truthfully measure the more relevant analytical impact on phylogenetic reconstruction.
- A quantitative evaluation of the position of cells in phylogenetic trees generated using ReDeeM filters -1 and -2 shows that they are mostly randomly repositioned with respect to each other (**Supplemental Fig. 2**). As such, the assertion that the two ReDeeM filtering methods are giving similar results is not supported upon statistical analyses via most recent common ancestor (MRCA) analyses (**Supplemental Fig. 2**, bottom; 33.3% is random).
- Evaluation of the k-NN analysis, which is repeatedly presented as a validation of the ReDeeM approach, shows that the methodological choices introduced in the original ReDeeM publication will produce the desired results on permuted and random data alike (**Supplemental Fig. 3**). Specifically, given the filtering for variants that co-segregate with the expected hematopoietic lineages (thereby excluding 70-80% of all variants from the downstream k-NN analysis), our reevaluation confirms that these metrics cannot present a form of validation of data or biology of any kind.

**Explanation for mtDNA mutational signature/enrichment profile**

- A noticeable omission from the ReDeeM method is a formal mixing experiment, either from species mixing (which is customary in single-cell technology development) or human donor mixing experiments (as we have previously performed for mtDNA analyses[4]). However, mixing experiments are critical to observe potential issues in library preparation and sequencing.

- For instance, ambient mtDNA molecules (cross-contaminating mtDNA that may occur during the joint lysis and transposition of cells as reported previously[4]) are not considered (**Supplemental Fig. 4**). Such variants would show the expected mtDNA mutational signature, but provide a source of error that cannot be corrected by consensus calling.
- Using Extended Data from the published manuscript that mixed cells of 6 human donors together[3], we performed genetic demultiplexing, doublet inference, and donor-specific homoplasmic (germline) variant calling (**Supplemental Fig. 5**). Specifically, we used a very lenient threshold for doublet classification (3,679 out of 10,745 barcodes with 10x mtDNA coverage) to establish a lower bound for the contamination. From these analyses (as we have previously used[4]), we estimate the contamination rate of ambient mtDNA to be 11.5% in ReDeeM, which will obscure phylogenetic relationships upon inclusion of mtDNA variants only supported at low molecular copy number.

**Conclusions**

Our reanalysis of original ReDeeM data with filters -1 and -2 demonstrates that the two filters yield vastly diverging results. The fact that the connectivity metrics and resulting phylogenetic trees are substantially different between the two filters only reinforces our original concern that artifactual mtDNA variants (as now removed by filter -2) remain a substantial driver of the purported phylogenetic signals. The repeatedly presented k-NN analysis is flawed by design and cannot be considered a validation of the ReDeeM methodology, nor does it provide support for the validity of artifactual variants. Additional confounders that affect the robustness of variants with single-molecule support for clonal and phylogenetic inference are not considered. The authors maintain that variants supported by only one molecule remain informative for phylogenetic inference by emphasizing observing the expected mtDNA mutational signature profile. However, our estimation of the contamination rate suggests that ambient mtDNA presents a significant confounder of the ReDeeM method. Notably, the contamination rate is substantially higher than previously reported for mtscATAC-seq[4], which will require further investigation but only support the notion that mtDNA variants with low molecule copy number support shall not be considered for phylogenetic inferences.

Finally, the results remain inconsistent with i) properties of mitochondrial genetics and its multi-copy-number concerning the faithful inheritance of variants supported by single molecules at low variant allele frequencies and ii) phylogenetic analysis of hematopoietic colonies based on whole genome sequencing, which directly compared the utility of mtDNA to gold-standard nuclear variants for deriving accurate phylogenies[5]. No attempt is made to reconcile these differences. Thus, we conclude that the findings reported in the original *Nature* publication[3] remain substantially flawed.

### Supplemental Figures

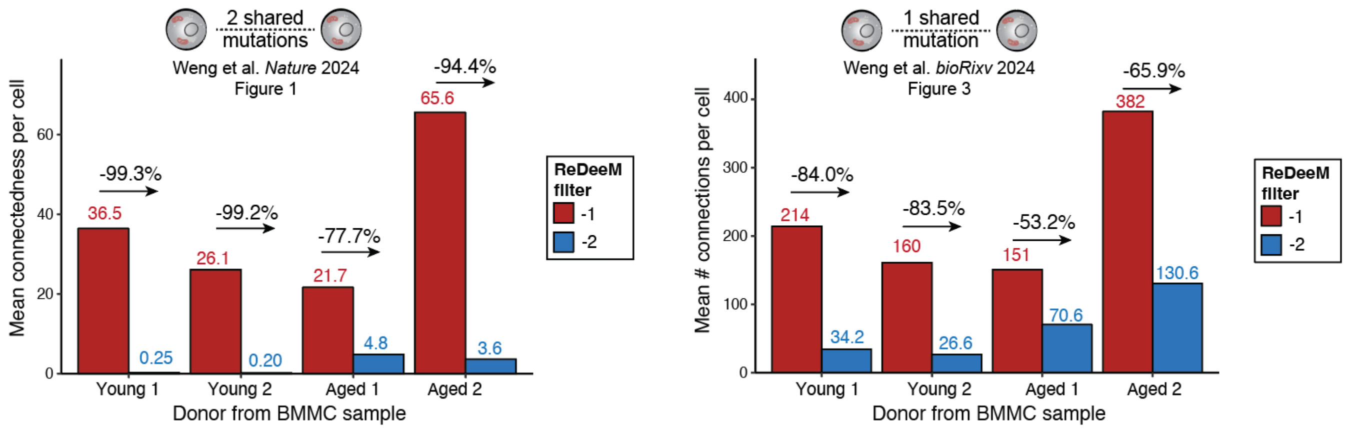


**Supplemental Figure 1. ReDeeM-based cell-cell connectivity is substantially reduced between filter choices -1 and - 2.** Schematic of key parameters indicating the number of shared mutations required to establish connectivity between two cells as previously applied by ReDeeM (top). The mean number of connections per cell (y-axis) is shown with the relative differences between ReDeeM filters -1 and -2 being indicated in % across bone marrow mononuclear cells (BMMCs) from four profiled donors (bottom).

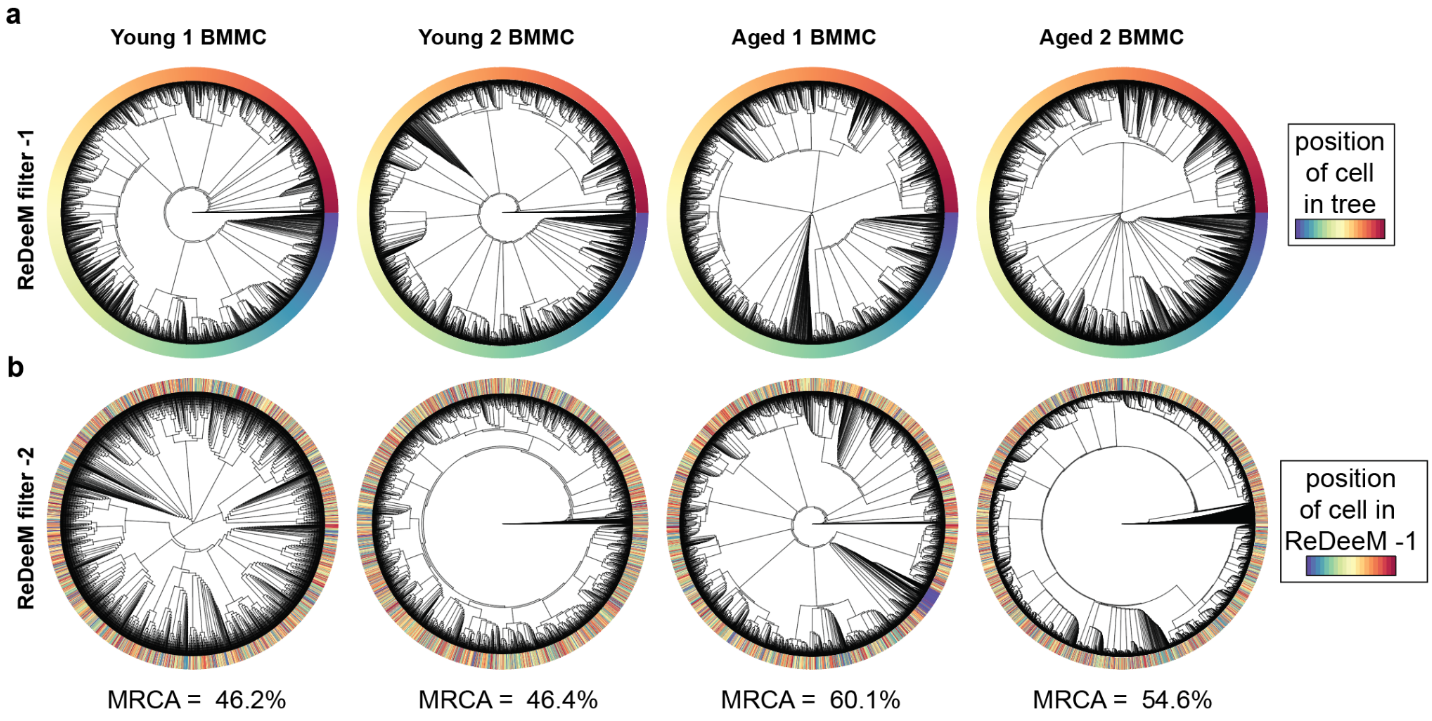


**Supplemental Figure 2. Inference of phylogenetic trees by ReDeeM filters -1 and -2 lead to vastly diverging distributions of hematopoietic cells.** Phylogenetic trees of BMMCs from four donors (columns) were constructed using (**a**) ReDeeM filters -1 and (**b**) -2, with filter -2 removing edge variants. The color bar marks the relative position of cells in the topology using ReDeeM filter -1 (top), showing the substantial reshuffling of cells after applying ReDeeM filter -2 (bottom). Most recent common ancestors (MRCA) are quantified in % for the lower tree using the top tree as a reference; 33.3% presents a random result; 100.0% presents a perfectly replicated tree.

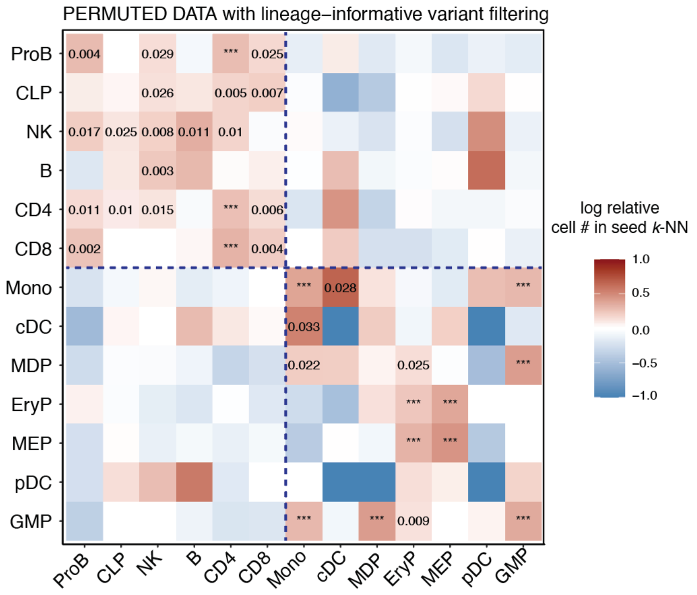


**Supplemental Figure 3. K nearest neighborhood (k-NN) analysis on permuted cell labels**. This analysis shows similar lineage enrichment results compared to the original data, presenting a major analytical flaw in the ReDeeM analysis workflow. Specifically, it pre-filters lineage-enriched mtDNA variants by requiring an enrichment in a known hematopoietic lineage[3] only after which the k-NN analysis is conducted. Compare to Fig. 4C, ReDeeM filter -2, all mtDNA mutations[1].

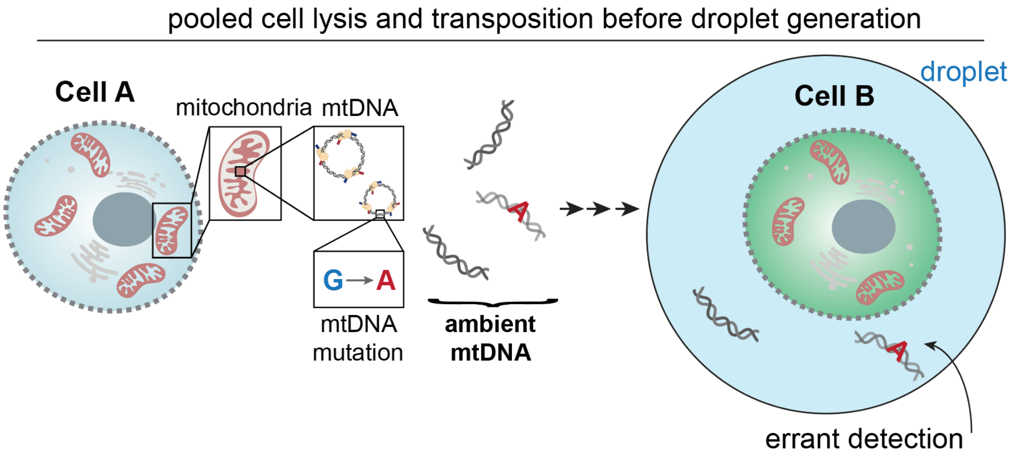


**Supplemental Figure 4. Ambient mtDNA molecules contribute to the cross-contamination of mtDNA variants in droplet single-cell workflows.** Like ambient mRNA[6,7], mtDNA molecules from cell A may be released during cell lysis and transposition in Multiome/ATAC-seq workflows (left). The mtDNA molecule may then be errantly droplet-encapsulated with cell B, leading to the cross-contamination of mtDNA molecules and variants between cells as previously demonstrated[4]. Such errors cannot be accounted for by either ReDeeM workflow and may further confound the utility of mtDNA variants only supported by single molecules for clonal and phylogenetic inference.

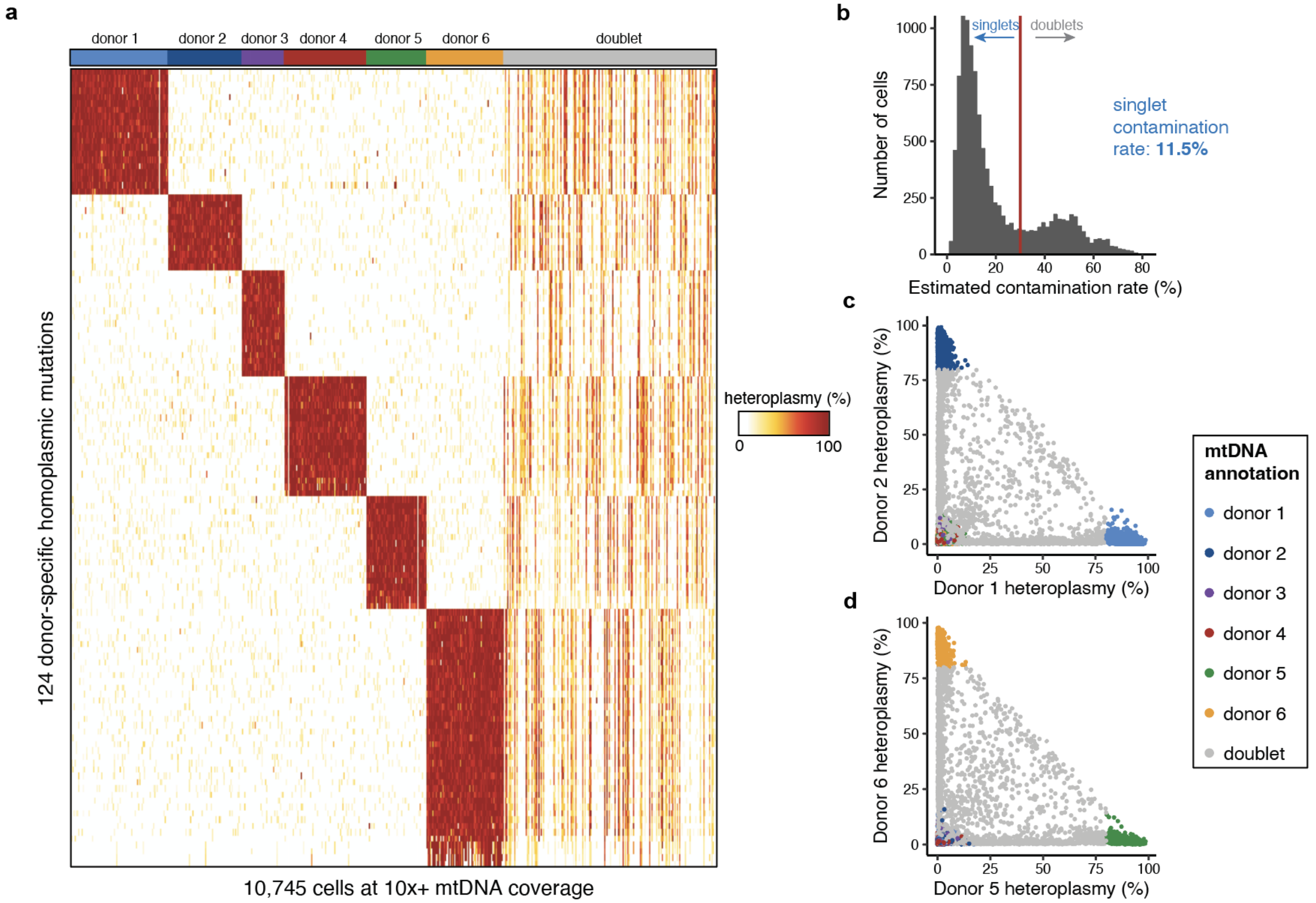


**Supplemental Figure 5. Quantification of cross-contamination of ambient mtDNA in the ReDeeM data.** **(a)** Cell-by-variant heteroplasmy matrix of 124 homoplasmic variants specific to one of the six donors that were jointly profiled in the extended donors mixing experiment[3] ("Young BMMC"). **(b)** Summary of the mean contamination rate of cross-donor homoplasmic variants per barcode, defined as the percent of heteroplasmy from donors excluding the top individual. The mean contamination rate per singlet is 11.5%. **(c)** Exemplary scatter plot of donor 1 and donor 2 mean heteroplasmy of otherwise donor-specific homoplasmic SNPs, colored by annotation (columns in (a)). **(d)** Same as in (c) but for donors 5 and 6.
